# ARID3a-Expressing Naïve B Cells in SLE have an Activated Phenotype and Transiently Express Surface CD68

**DOI:** 10.64898/2026.07.10.737835

**Authors:** Joshua Garton, James Hocker, Lori Garman, Hua Zhong, Kurt Zimmerman, Joel M. Guthridge, Judith A. James, Carol F. Webb

## Abstract

Numbers of ARID3a (<u>AT</u>-<u>R</u>ich Interaction <u>D</u>omain <u>3a</u>) -expressing B lymphocytes from patients with systemic lupus erythematosus (SLE) are associated with increased disease activity. Normally, ARID3a-expressing circulating naïve B cells are rare, but in SLE naïve B cells dramatically increase ARID3a expression. We found that *in vitro* stimulation of B lymphocytes from healthy individuals with a cocktail of cytokines and agonists induced ARID3a in a subset of activated naïve B cells and in IgD^-^CD27^-^ double negative B cells previously associated with autoimmunity. Single cell RNA-seq of isolated naïve B cells from ten SLE patients, with varying frequencies of ARID3a-expressing cells, revealed that ARID3a-associated genes included activation markers. Moreover, our data revealed the unexpected co-expression of the scavenger receptor CD68 with ARID3a, at both the transcript and protein level, in activated subsets of naïve B cells. Inhibition of ARID3a in stimulated B cell cultures blocked naïve B cell activation and CD68 expression. Together, these data identify ARID3a and CD68 as markers of naïve B cell precursors associated with autoimmunity in SLE.

## INTRODUCTION

Systemic lupus erythematosus (SLE) is a devastating and heterogenous autoimmune disease characterized by the loss of immunological tolerance and production of autoantibodies. The mechanisms responsible for tolerance breaches are unknown but result in increased populations of autoantibody-producing B lymphocytes not found in healthy individuals. Double negative (DN) B cells that lack surface proteins CD27 and IgD are enriched in SLE patients with active disease and nephritis and are produced by maturation of a subset of “activated” naïve B cells ^1–3^. These activated naïve B cells differ from healthy control naïve B cells in chromatin accessibility signatures ^4^, suggesting that early activation events at the naïve B cell stage contribute to SLE pathogenesis.

ARID3a (A+T-rich interaction domain 3a) is a DNA-binding protein originally discovered as a component of a protein complex that upregulates immunoglobulin heavy chain transcription and is associated with transcription enhancing and suppressing functions in a cell type-specific fashion ^5–9^. Generation of transgenic mice with constitutive expression of ARID3a in all CD19^+^ B cells was sufficient to generate autoimmunity with production of anti-nuclear antibodies and glomerulonephritis ^10,11^. Likewise, increased disease activity in SLE patients is associated with increased numbers of ARID3a-expressing B lymphocytes ^12^. Longitudinal studies of SLE patients showed that numbers of ARID3a^+^ B cells in individual patients varies over time, and that ARID3a can be expressed in all stages of B cell differentiation, including naïve B lymphocytes, a developmental stage where ARID3a expression is rare in healthy controls ^12^.

Although we cannot readily determine what signals induced ARID3a expression in individual SLE patients, others demonstrated that Epstein Barr Virus (EBV) both induces and requires endogenous ARID3a expression for viral EBNA-1 protein production ^13^. We and others have linked recurrence of EBV with autoimmune responses in SLE ^14–16^. We hypothesized that expression of ARID3a in naïve B cells in SLE patients might contribute to gene expression patterns associated with autoimmunity and that the ARID3a^+^ naïve B cell population might contain precursors of autoantibody-producing cells. Our previous data support the idea that ARID3a expression is associated with cell-type specific differences in gene expression ^17,18^. Therefore, we performed *in vitro* experiments to induce activated naïve and DN B cells to determine if ARID3a is induced in this model system. In addition, we performed single cell RNA-seq (scRNAseq) experiments on isolated SLE naïve B cells to define the transcriptome of this subset of ARID3a-expressing cells that contains the “activated” naïve B cells proposed by others to be precursors of pathogenic autoimmune B cells ^1^.

## RESULTS

### ARID3a expression in naïve B cells can be induced *in vitro*

Others found that activated subsets of naïve B cells are precursors of pathogenic DN B cells in SLE and that similar types of activated naïve B cells could be induced *in vitro* from healthy control B cells with a complex milieu of stimulants ^1^. Fig. 1 shows a representative *in vitro* experiment in which isolated total B cells from a healthy control were stimulated with the TLR agonist R848, anti-IgM F(ab’)2, and growth factors BAFF, IL21, IL-2 and IFNγ for three days and were assessed for DN and naïve B cell subsets with ARID3a by flow cytometry. The majority of “activated” naïve (IgD^+^CD27^-^) B cells, but not the CD21^+^ “resting” naïve B cells from the same culture, expressed ARID3a. Similarly, the more mature IgD^-^CD27^-^ cells that maintain CD21 expression did not express ARID3a, while the DN2 cells that had lost CD21 expression were predominantly ARID3a^+^ by 3 days of culture, a time point before we observed expression of CD11c. This stimulation model is the first system in which ARID3a expression has been induced in naïve B cells *in vitro*. In addition, these data indicate that ARID3a is expressed in activated naïve B cells, the proposed precursors of pathogenic DN B cells, many of which also express ARID3a.

**Figure 1.**
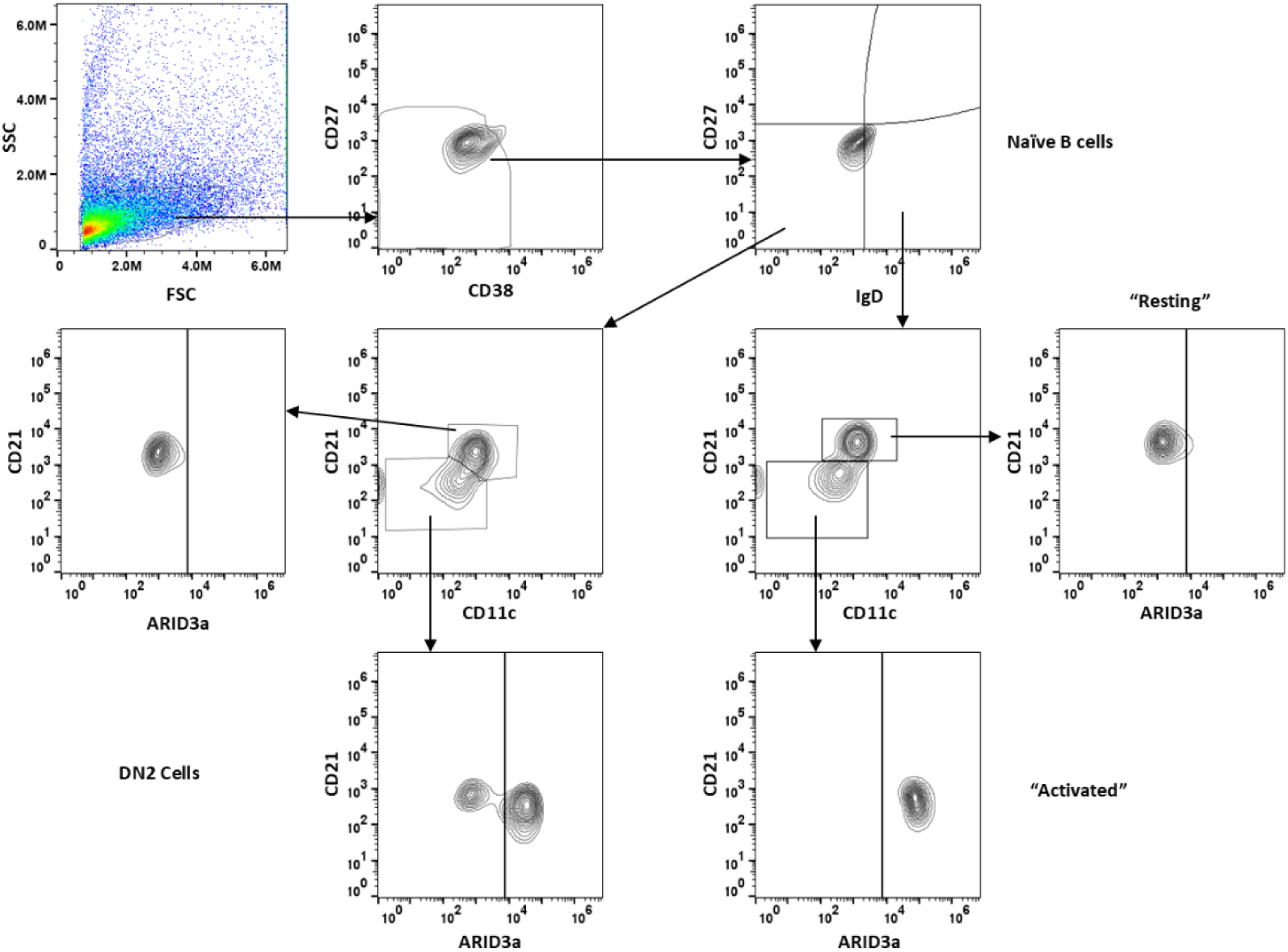
“Activated” naïve B cells and DN2 cells express ARID3a. Healthy donor B cells enriched by magnetic bead depletion of other immune cells and were stimulated with 1 µg/ml R848, 10 µg/ml F(ab’)2 anti-IgM, 10 ng/ml each BAFF and IL21, 50 U/ml IL-2 and 20 ng/ml IFNϒ for 3 days prior to analyses by flow cytometry using the markers and gates shown. Representative gating of DN2, activated, and resting B cells is shown. N=8.

### ARID3a-expressing naïve B cells express distinct transcriptome

Our previous data showed both ARID3a^+^ and ARID3a^-^ naïve B cells, at varying frequencies, occur in individual SLE patient peripheral blood samples ^12^. Therefore, we used single cell RNA-sequencing (scRNA-seq) to define and identify differentially expressed genes (DEGs) associated with ARID3a expression within the naïve B cell population in SLE. We first confirmed differential ARID3a transcript expression within total SLE B cells (SFig. 1A) and detection of ARID3a transcript within flow cytometrically sorted naïve B cells (SFig. 1B). We observed strong correlations between ARID3a protein and transcript levels in naïve B cells, unlike our observations with other blood cell types which can express ARID3a transcript without protein ^17^. FACS-sorted naïve B lymphocytes (IgD^+^IgM^+^CD10^-^CD27^-^) were isolated from ten randomly selected SLE patients with a wide range of disease activity. CD10 was used as a marker to exclude transitional B cells ^19^ which can retain ARID3a expression in mice ^20^. Unsupervised UMAP clustering of scRNA-seq data from all cells revealed five clusters (Fig 2A) and ARID3a^+^ cells were highly enriched in Cluster 2 (Fig. 2B). The top 20 DEGs expressed with each of the five clusters are shown in the heatmap in Fig. 2C. Using Partek software, we defined 134 ARID3a^+^ and 695 ARID3a^-^ cells based on >0.1 ARID3a transcripts per million (TPM) shown by violin plots of ARID3a transcript levels, in Log2(TPM) in each cell for each of the 10 patients (Fig. 2D) and revealing the heterogeneous nature of ARID3a expression among the ten individuals. ANOVA analyses identified 4,333 DEGs of 24,002 genes (FDR < 0.05, fold-change > +/- 2) in ARID3a^+^ vs. ARID3a^-^ cells as shown in the Volcano plot (Fig. 2E). Equivalent numbers of genes were upregulated (2185) versus down regulated (2148) with ARID3a. These data indicate that ARID3a^+^ naïve B cells are defined by distinct transcriptomes compared to naïve B cells without ARID3a.

**Figure 2.**
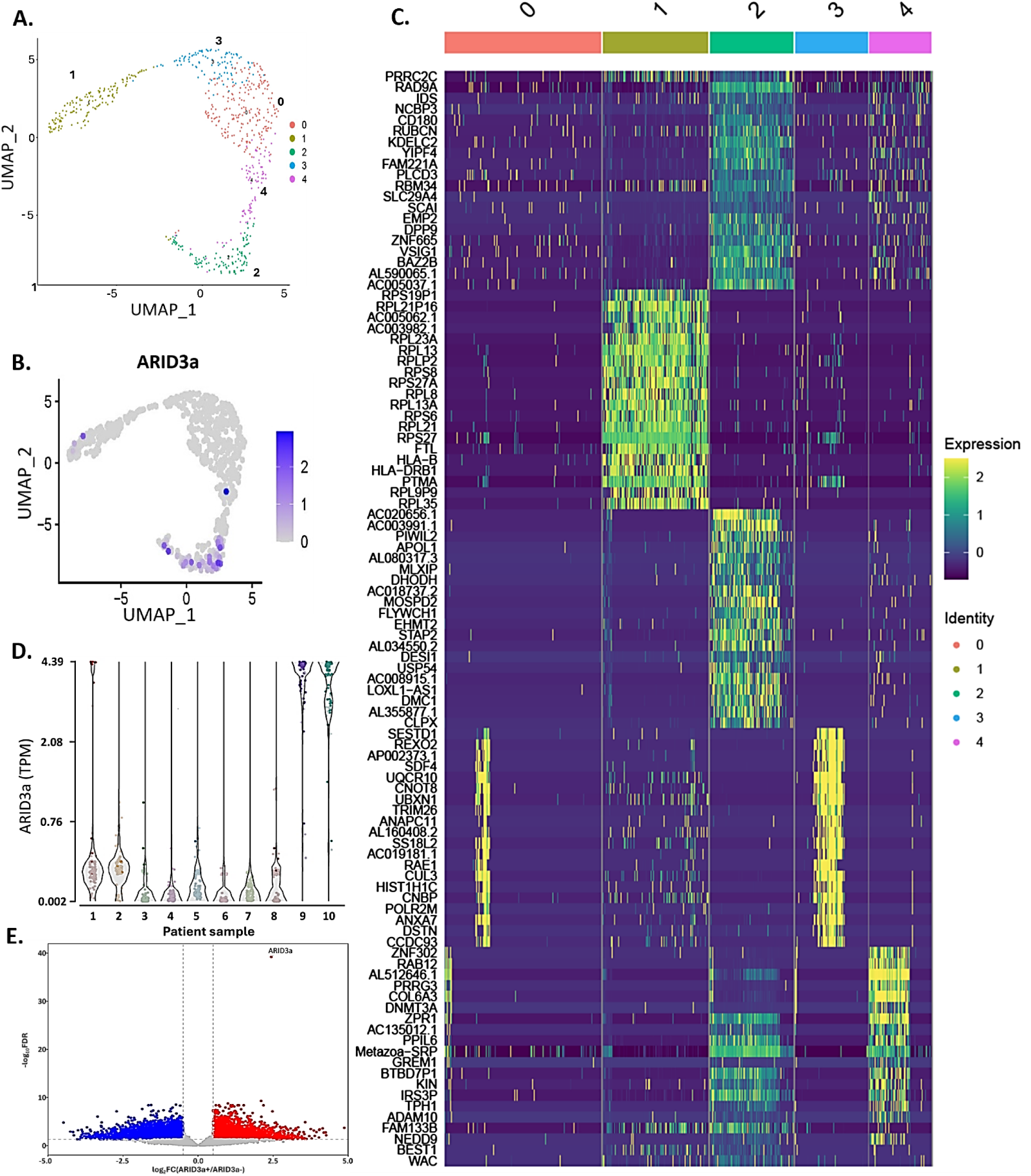
ARID3a^+^ SLE naïve B cells have a discrete transcriptome compared to ARID3a^-^ naïve B cells. Single cell RNAseq was performed with sorted naïve B cells (CD19^+^CD27^-^ IgD^+^CD10^-^) from 10 SLE patients. (A) Unsupervised cluster analyses revealed five groups. (B) ARID3a^+^ cells are depicted in purple and clustered largely into group 2 with a few cells in groups 1 and 4. (C) A heatmap of the twenty, most highly differentially expressed genes within each of the 5 groups is shown. (D) Violin plots of naïve B cells (individual points) from each SLE patient (samples 1-10) show ARID3a transcripts per million (TPM). (E) Volcano plot of DEGs in ARID3a^+^ vs ARID3a^-^ SLE naïve B cells. Each dot represents a gene, with blue indicating down-regulation, red indicating upregulation and grey indicating no significant change. The ARID3a gene is the uppermost point.

Several genes of interest, including *TLR7* the viral RNA sensor, and the activation marker, *PTK2,* were predominantly expressed in cells in cluster 2 that was enriched for ARID3a-expressing cells (Fig. 3A). Other genes associated with production of differentiation into DN2 cells ^1^, such as *BCL2, IL10RA, CXCR5, and HLA-DR* were not associated with cluster 2, ARID3a-enriched cells, but were more highly expressed in cluster 1 and/or other groups (Fig. 3B). BACH2 was previously shown to be expressed in resting naïve B cells ^1^. The transcript encoding the inhibitory receptor CD22 was highest in cluster 2 with ARID3a-expressing cells but was also abundant in cluster 1 and was present in all other clusters. Likewise, ICOSLG and CD200 ^21^ were highly expressed in cluster 2 but were also found in other clusters as well. These regulatory surface markers are consistent with cell activation.

**Figure 3.**
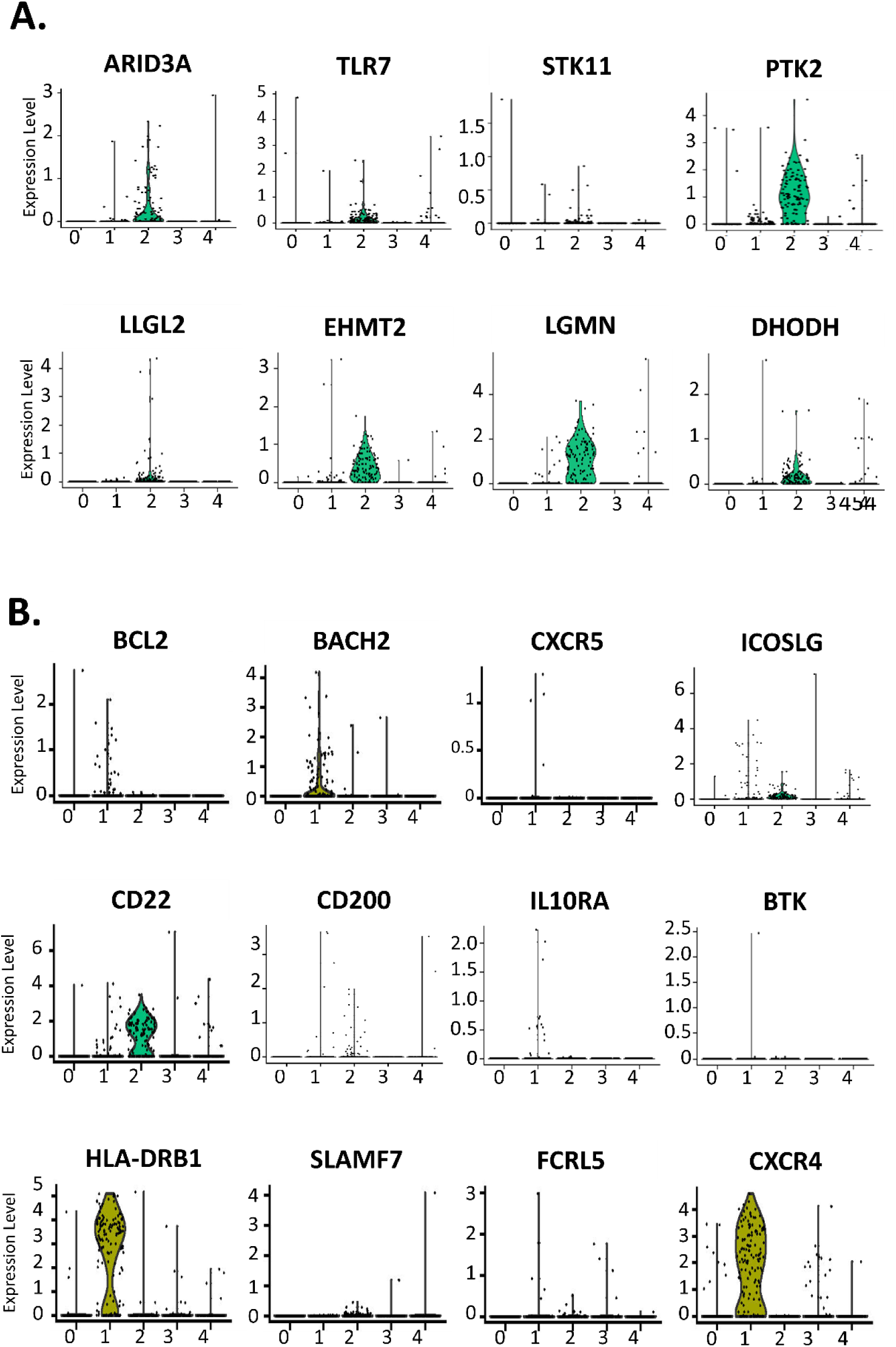
ARID3a expression is associated with multiple signaling and activation genes. Individual gene expression patterns are shown for each of the five clusters identified in Fig. 2A. (A) Violin plots show individual cells expressing ARID3a in comparison to other transcripts enriched in group 2 ARID3a-associated cells. (B) Violin plots of other lupus-associated genes did not group predominantly with ARID3a-associated cells.

### Psuedotime analyses suggest that cells expressing ARID3a are less differentiated than some ARID3a^-^ clusters

Psuedotime analyses were performed for the five clusters identified in Fig. 2a and a pseudotime plot and trace plot are shown (Fig. 4A and 4B). Fig. 4C shows the developmental potential of cells within each cluster, again suggesting at least a portion of the ARID3a-expressing naïve B cells are less differentiated than the majority of cells in cluster 2 which shows high expression of HLA markers, for example. The top 100 pseudotime significant genes presented via unsupervised clustering (Fig. 4D) emphasize differences between expression levels in ARID3a^+^ cells in cluster 2 and the more differentiated ARID3a^-^ naïve B cells in cluster 1. Although we cannot predict from these analyses which, if any, of the ARID3a^-^ clusters may be precursors of ARID3a^+^ naïve B cells, the data are consistent with transient expression of genes as naïve B cells become activated to differentiate.

**Figure 4.**
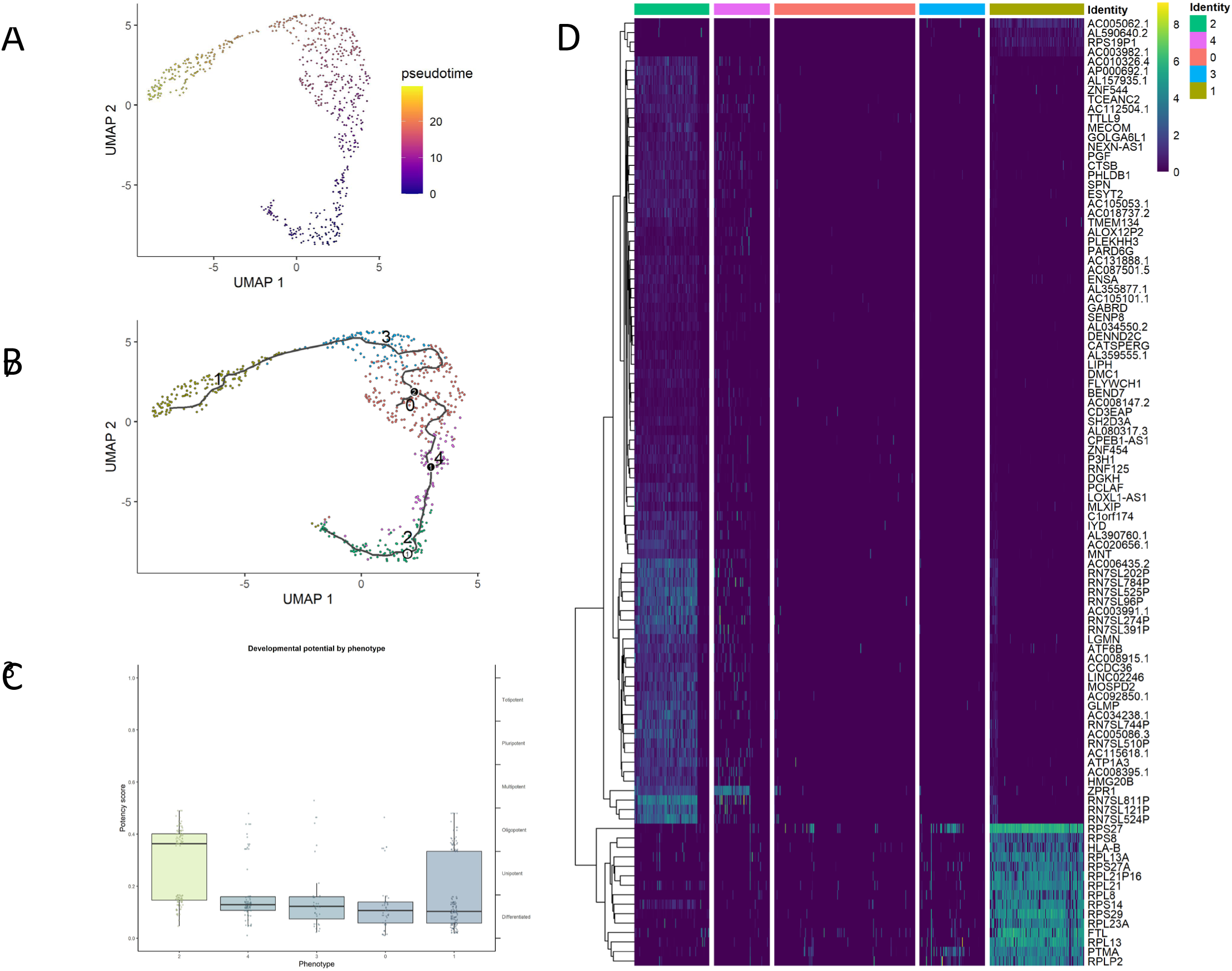
Pseudotime analyses indicate ARID3a^+^ naïve B cells are not fully differentiated. Pseudotime analyses of all naïve B cells were performed and are shown as a pseudotime plot (A) and trace plot (B). The developmental potential of each cluster was analyzed (C). Unsupervised clustering of the top 100 pseudotime significantly differentially expressed genes (p>2) are presented (D).

### CD68 is transiently expressed in activated ARID3a^+^ naïve B cells generated *in vitro*

ARID3a is an intracellular protein and no surface markers exist to readily distinguish ARID3a^+^ from ARID3a^-^ naïve B cells. Therefore, we searched for transcripts of surface proteins that might be strongly associated with ARID3a expression in activated naïve B cells. Several surface markers were upregulated in ARID3a^+^ naïve B cells, including CD22 (Fig. 3B), but were not limited to only ARID3a^+^ naïve B cells. However, both CD68 and CD36 transcripts were significantly expressed within cluster 2 ARID3a-associated cells. While the significance of increased CD36 expression was skewed by increased expression levels of a few cells, CD68 expression was abundant in ARID3a-associated cells (Fig. 5A). Fluorescence staining of *in vitro* stimulated healthy naïve B cells containing activated ARID3a^+^ cells confirmed that a subpopulation of ARID3a^+^CD68^+^ cells was present (Fig 5B). Gating on those CD68^+^ARID3a^+^ cells revealed that those cells were naïve IgD^+^CD27^-^ B cells, while the ARID3a^-^ cells were largely CD27^+^ (Fig. 5B). ARID3a^+^CD68^-^ cells had a DN phenotype. As expected, total B cells stained with ARID3a and CD36 did not show CD36 protein within ARID3a-expressing groups of cells (Fig. 5C). These data suggest CD68 is a marker for activated naïve B cells induced *in vitro*.

**Figure 5.**
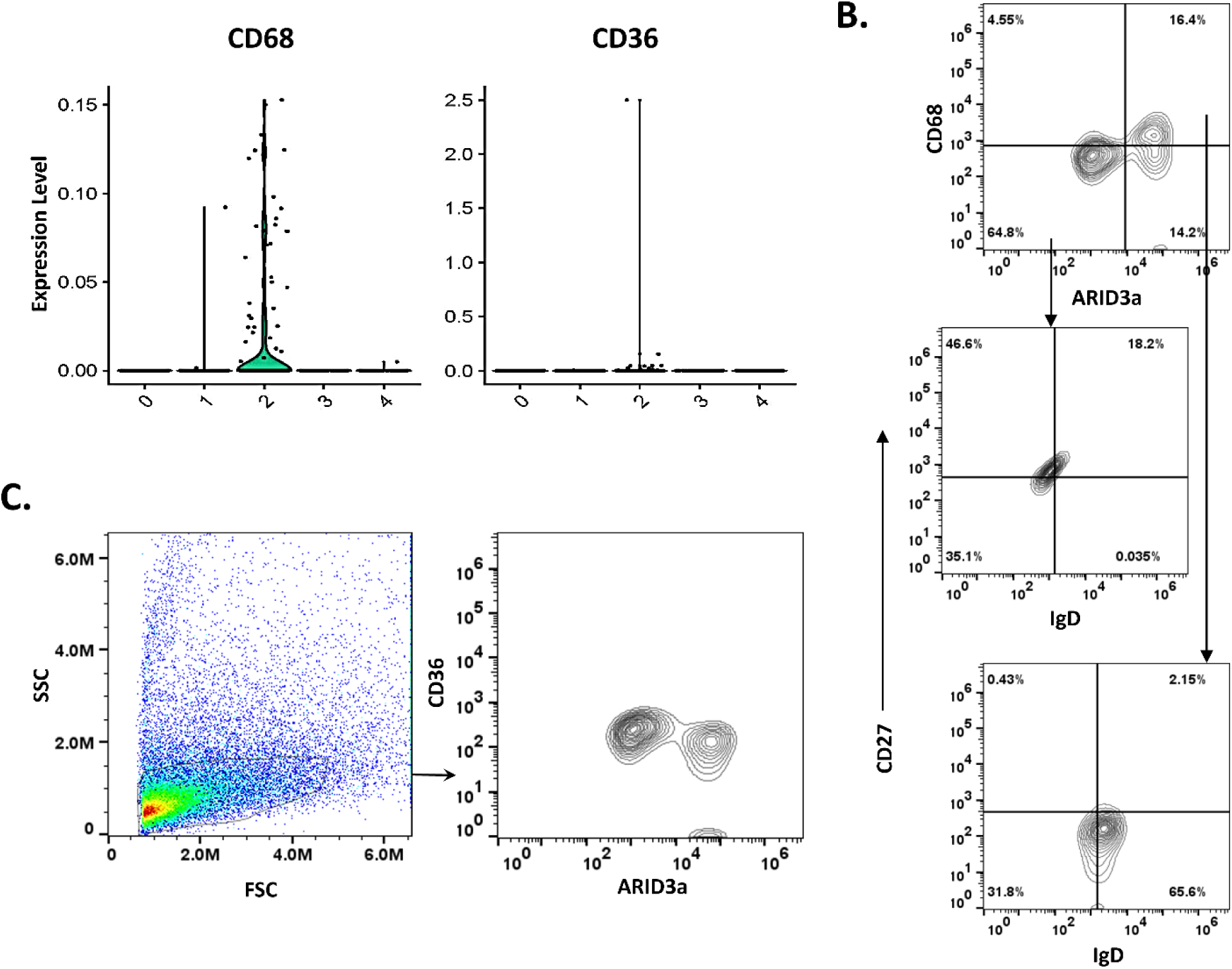
CD68 is a surface marker associated with ARID3^+^ activated naïve B cells. (A) Violin plots from scRNA-seq data show numbers of cells expressing the surface markers CD68 and CD36 in each cluster group. (B) Flow cytometric analyses of total healthy control B cells stimulated in culture for three days, as in Fig. 1, show ARID3a and CD68 expression. Further gating for ARID3a^+^CD68^+^ cells reveals that these are largely naïve B cells expressing IgD and lacking CD27. (C) The same cells were stained for ARID3a and CD36 and show live gating. Quadrants were set by isotype controls. N=5.

### Activated naïve B cells from SLE patients co-express ARID3a and CD68

We also examined B cell subsets directly *ex vivo* from peripheral blood of healthy controls and SLE patients and found that while healthy control naïve B cells did not express ARID3a or CD68 (Fig. 6A), ARID3a^+^ naïve B cells from SLE patients also co-expressed CD68 (Fig. 6B). However, SLE patient samples with no ARID3a^+^ naïve B cells also failed to express CD68 (Fig. 6B). These data are consistent with our previous data showing ARID3a-expressing naïve B cells were not detectable in all SLE patients (Fig. 2D and ^12,22^). In this data set, at a single time point, 60% of randomly assessed SLE patients had ARID3a^+^CD68^+^ naïve B cells compared to our previous findings of 63% of patients with increased numbers of naïve B cells expressing ARID3a ^12^. Comparisons of tSNE plots show locations of gated B cell subsets within the entire pBMC population and demonstrate the absence of activated naïve B cells in healthy controls (Fig. 6C). Together, these data suggest that CD68 is transiently expressed with ARID3a on activated naïve B cells.

**Figure 6.**
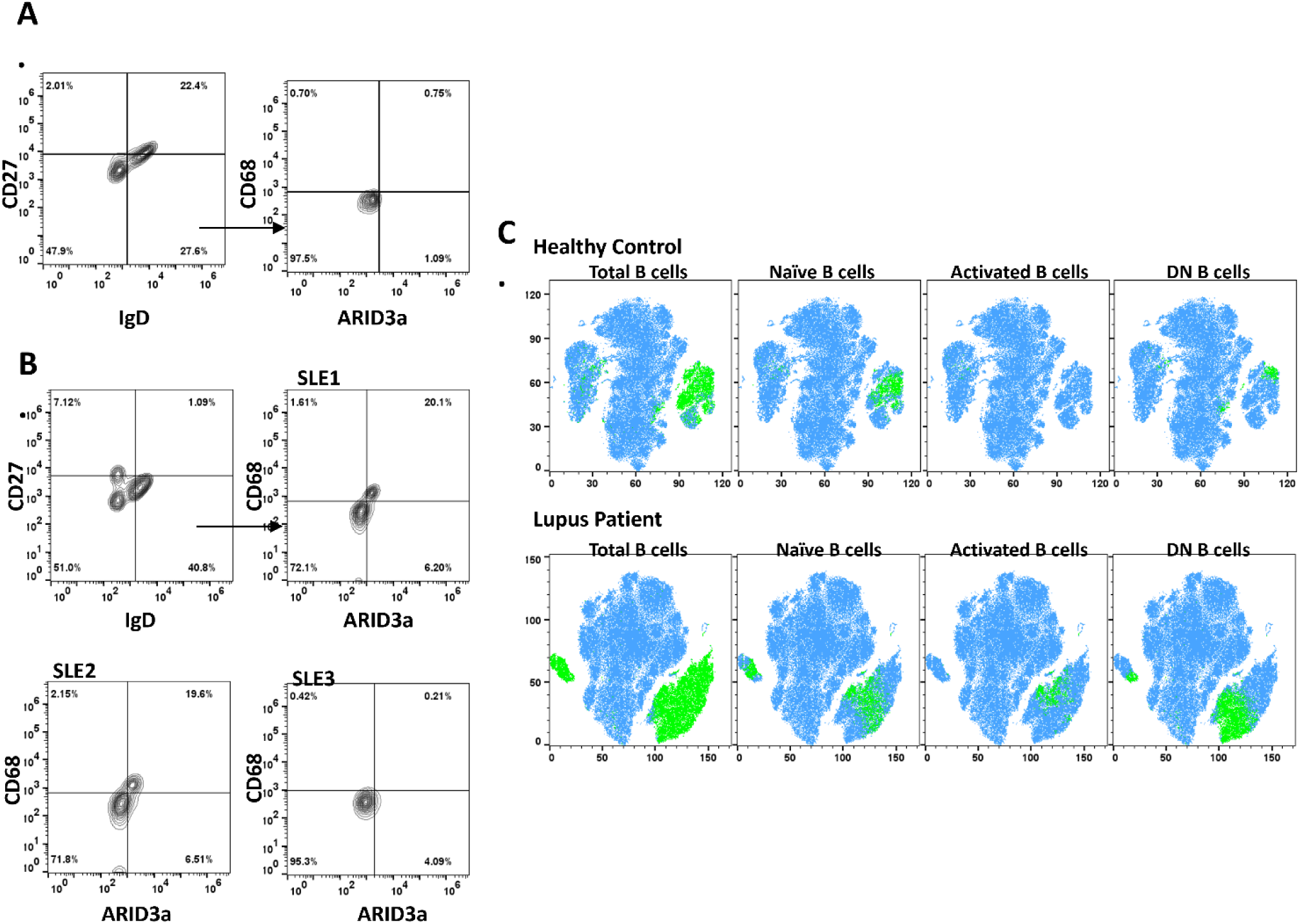
CD68 is co-expressed with ARID3a in naïve B cells from SLE patients. (A) Flow cytometric analyses of CD19^+^ B cells from one representative healthy control (N=6) and (B) three SLE patients show CD68 expression in association with ARID3a (N=20). Quadrants were set as described. (C) Representative spectral flow cytometry tSNE plots demonstrate location of “activated” naïve ARID3a^+^ CD68^+^ and DN B cells within the total CD19^+^ B cell subset of a healthy control and an SLE patient.

### ARID3a inhibition reduces activation of naïve B cells *in vitro*

To determine if ARID3a expression is important for *in vitro* induction and differentiation of B cells, perhaps into DN B cells as suggested by Jenks, et al. ^1^, we stimulated isolated healthy B cells, as in Fig. 1, both with and without vivo-morpholinos specifically targeting ARID3a ^23^. We observed that inhibition of ARID3a at the initiation of culture dramatically inhibited cell expansion on day 3 (Fig. 7A) as evident by the absence of expanded clumps of blast-like cells. By 3.5 days, control cultures and cultures treated with an unrelated morpholino contained activated naïve B cells expressing ARID3a and CD68 while cultures treated with ARID3a-specific morpholinos showed reduced ARID3a^+^CD68^+^ cells (Fig. 7B). Consistent with our previous findings and reports by others suggesting that vivo-morpholinos remain active for a period of 3-4 days *in vitro* ^24^, we observed redevelopment of clumps again by day 5 of culture without additional vivo-morpholino (not shown). ARID3a inhibition resulted in significant reduction numbers of IgD^+^ naïve B cells compared to stimulated control cultures (Fig. 7C). In addition, percentages of B cells expressing the activation marker CD27 were also significantly reduced (Fig. 7D), as were the activation markers CD11c, HLA-DR and CD38 (Fig. 7E). These data suggest that ARID3a actively contributes to production and/or expansion of CD68^+^ activated naïve B cells.

**Figure 7.**
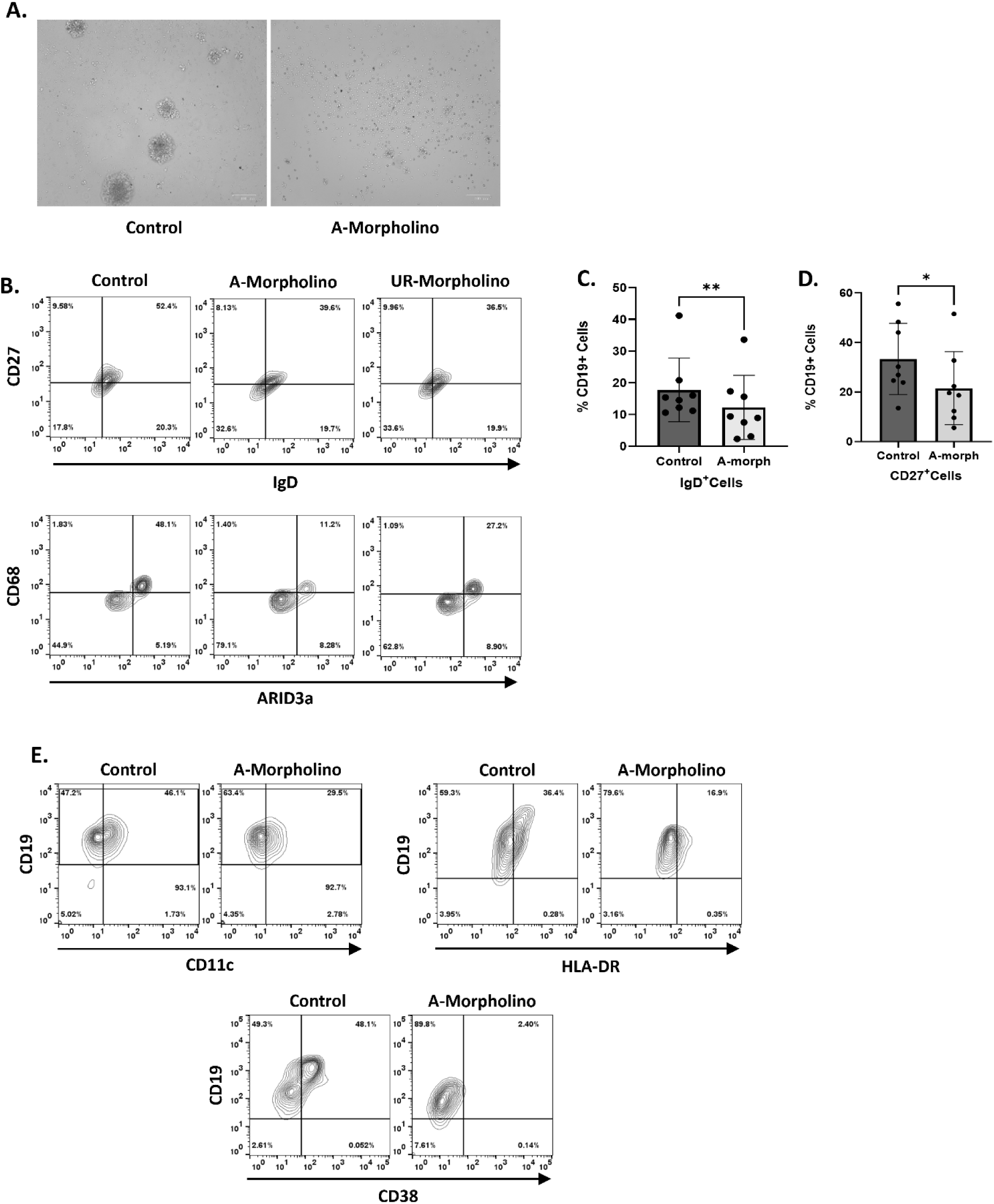
ARID3a is required for B cell activation *in vitro*. (A) Healthy donor B cells enriched by magnetic bead depletion were stimulated as in Fig. 1 and cultured for 3 days with or without 5 µg/ml ARID3a-specific morpholino. (B) Total CD19^+^ B cells cultured 4 days with and without ARID3a-specific (A) or an unrelated control (UR) morpholino were gated for naïve IgD^+^CD27^-^ cells (top panels) and ARID3a and CD68 (bottom panels). (N=6) (C) Percentages of cells expressing IgD and CD27 (D) without and with A-morpholino treatments for 3-5 days were assessed by flow cytometry (N=8, **p= 0.0097 and *p=0.0253 respectively, by two tailed t-tests). (E) Flow cytometry shows inhibition of activation markers CD11c, CD38 and HLA-DR in ARID3a-morpholino treated cultures versus control cultures (N=3).

## Discussion

Data in this study show that ARID3a is expressed in activated naïve B cells. Activated naïve B cells are reported to contain precursors of DN2 autoreactive B cells that are pathogenic in autoimmune diseases such as SLE ^1^. Using scRNA-seq of isolated naïve B cells from SLE patients, we identified multiple genes associated with ARID3a expression in naïve B cells. Moreover, we identified CD68 as an unexpected biomarker for activated, ARID3a^+^ naïve B cells in both stimulated healthy B cell cultures and within activated naïve B cells in SLE blood samples. We observed that CD68 expression was not maintained in cells that developed into DN B cells. Together, these data demonstrate the potential importance of ARID3a expression in activated naïve B cells and support our previous data associating numbers of ARID3a-expressing B cells with increased disease activity in SLE ^12^.

Both transcript and protein analyses identified the scavenger receptor CD68 as a novel surface marker co-expressed with ARID3a on activated naïve B cell subsets. CD68 is typically known as a monocyte marker; however, earlier studies also reported that select anti-CD68 monoclonal antibodies stained some human lymphocytes ^25^. As in that study, we noted variability in the specificity of staining of individual anti-CD68 monoclonal antibodies to the ARID3a^+^ naïve B cells, perhaps indicating a specific motif or post-translational modification is significant for binding. We speculate that CD68 expression in SLE ARID3a^+^ naïve B cells may have been missed by some 10x studies which may have dismissed small subsets of cells with two lineage markers, particularly if those cells are not present in healthy controls or all lupus samples. Numbers of ARID3a^+^ naive B cells are highly variable in SLE patients and within individuals over time and only about two-thirds of our randomly collected SLE samples showed increased numbers of naïve ARID3a^+^ B cells ^12^. Although ARID3a is a low abundance transcript, data mining of naive B cell transcripts from another study ^1^ also identified low levels of CD68 and ARID3a within SLE samples. CD68 was also found on a subset of mouse marginal zone B cells ^26^, a population of B cells also associated with autoimmunity ^27^. Additional experiments will be required to determine if the naïve B cells that co-express ARID3a and CD68 in human cells are marginal zone B cell precursors or B1-like lineage cells, another B cell subset associated with both autoimmunity and ARID3a expression in mice ^20,26,28,29^. Alternatively, transient CD68 expression could be a normal developmental marker of activation in all conventional naïve B cells. Regardless, CD68 is a marker of activated naïve B cells that can be used to isolate cells with and without ARID3a for future functional studies.

ARID3a modulates gene expression, and many genes showed upregulated expression in ARID3a^+^ naïve B cells compared to resting naïve B cells. CD22, a marker of marginal zone precursors^30^, is expressed in ARID3a^+^ naïve B cells (Fig. 3B) and in pathogenic DN2 cells^1^, while other DN2 cell activation markers, such as HLA-DR and IL10RA were primarily found in naïve B cell groups without ARID3a (Fig. 3). Likewise, highly expressed genes in DN1 cells such as BCL2 and CXCR5 were not co-expressed with ARID3a. However, TLR7 expression was abundant in ARID3a-associated cells and is a necessary mediator for DN2 cell induction ^1^. One of the most enriched genes associated with ARID3a-expressing cells is PTK2/FAK which is phosphorylated by BCR signaling and contributes to cellular activation^31^. EHMT2, a histone methyl-transferase, may contribute to alterations in gene expression at the chromatin level. Further analyses will be required to identify genes directly modulated by ARID3a. However, it is unlikely that ARID3a directly regulates CD68 expression because more mature DN ARID3a-expressing cells do not co-express CD68, and both CD68 and ARID3a can be expressed without the other protein in other cell types. Further exploration of the roles of ARID3a-associated genes and their functions in the production of autoimmune cells is important.

EBV requires the induction of endogenous ARID3a for the expression of viral EBNA proteins ^13^. Reactivation of EBV has been implicated as a predisposing event for lupus flares^14,15,32^. Antibodies to EBNA proteins have been associated with autoimmunity in both SLE and multiple sclerosis (MS) ^16,33^. Our preliminary data show the presence of ARID3a and CD68 co-expressing naïve B cells in subsets of MS patients and some patients with other types of autoimmune disease. SLE is a waxing and waning disease, and numbers of ARID3a-expressing B cells also vary within the naïve B cell subset (Fig. 5 and ^12,22^), perhaps providing a future tool for screening for upcoming increases in autoimmune responses seen in flares of disease. In addition, targeting these cells may have future therapeutic potential.

Data limitations include the fact that healthy, activated naïve B cells induced *in vitro* may not be identical to the ARID3a^+^ naïve B cells present in SLE patients. Others found that anergic naïve B cells lack CD21^34^ and have down-regulated Ig surface receptors making up 25-50% of SLE naïve B cells and 5-20% of healthy control naïve B cells ^35,36^. Therefore, such anergic B cells may account for the “activated” naïve B cells expanded from healthy controls after stimulation *in vitro*. In addition, our data do not exclude the possibility that other subsets of ARID3a^-^ cells may also contribute to pathogenic autoantibody-producing cells. For example, memory B cells also contribute to autoimmunity^37^. Nonetheless, these data help define an important subset of activated naïve B cells that may contribute to future autoimmune responses.

## METHODS

### Study design

The goal of this study was to define characteristics of ARID3a-expressing naive B cells that we previously identified in lupus patients compared to healthy individuals who do not have ARID3a-expressing naïve B cells. We used established, *in vitro*-stimulation assays to activate purified healthy B cells and generate cells resembling the pathogenic B cells found in SLE patients. ARID3a expression was induced in a subset of activated naïve B cells. scRNA-seq of cytometrically isolated naïve B cells from randomly selected SLE patients was used to identify transcriptomes of ARID3a-expressing naïve B cells and to search for potential surface markers limited to those activated B cells. CD68 was identified as an enriched transcript and a transiently expressed surface protein of ARID3a^+^ naive B cells in SLE blood and as a marker of activated ARID3a^+^ naive B cells induced *in vitro* from healthy individuals.

### Patient and healthy samples

Peripheral blood mononuclear cells (PBMCs) from individuals with SLE were obtained after consent from the Oklahoma Medical Research Foundation (OMRF) under IRB# 6634 during normal clinic visits. Age and sex-matched healthy control samples were either obtained from the OMRF repository or were purchased from the Oklahoma Blood Institute. Demographics are listed in Table 1.

**Table 1.** Human Demographics for PBMC Samples.

| **Groups** | **Age Range (Mean)**<br>years | **Sex**<br>F /M | **Ethnicity**<br>C / AA / NA / O |
| --- | --- | --- | --- |
| SLE patients | 22 - 82 (45.4) | 29 /4 | 17/ 11 / 3 / 2 |
| Healthy controls | 28 - 71 (54.9) | 10 /4 | 9/ 5/ 0 / 0 |

Caucasian (C); African American (AA); Native American (NA); more than one race (O)

### Cell isolation and stimulation

SLE and healthy control PBMCs were isolated from blood samples as previously described ^22^, and used immediately for flow cytometry or cryopreserved for later use. Briefly, PBMCs were thawed into 10mL of RPMI + 5% FCS (and 1µL benzonase), centrifuged at 1000 rpm for 7 mins and the cell pellet washed twice with 10 mL RPMI + 5% FCS. For scRNA-seq, human CD19^+^ cells were positively selected from SLE PBMCs with the REAlease CD19 MicroBead Kit (Miltenyi Biotec Cat# 130-117-034). Naïve B cells (CD27^-^, CD10^-^, IgD^+^) were then sorted using a BD FACSAria Fusion (BD Biosciences) prior to use. Healthy control B cells were subjected to magnetic bead purification via negative selection using an Easysep human B cell enrichment kit (Stemcell Technologies, #19054) and were resuspended at 10^5^ cells/ml in RPMI1640 with 10% FBS and were stimulated for three to five days with 1 µg/ml ml R848, 10 µg/ml F(ab’)2 anti-IgM (Invitrogen Cat# A24490), 10 ng/ml BAFF (BioLegend Cat# 559602) and IL21 (BioLegend Cat# 571202), 50 U/ml IL-2 (BioLegend Cat# 589102) and 20 ng/ml IFNγ (BioLegend Cat# 570204), with and without 5 µM ARID3a-specific vivo-morpholinos, as described ^1,23^.

### Antibodies and reagents used

Antibodies, cytokines and reagents used in this study are listed in Table S1.

### Flow cytometry

PBMCs were stained for 20 min on ice with various antibody combinations against cell surface markers after washing twice in FACS buffer. Following surface marker staining, cells were washed in FACS buffer, fixed with fixation buffer (BioLegend Cat # 420801), permeabilized with FoxP3/Transcription Factor Staining Buffer Set (Invitrogen eBioscience Cat # 00552300) and stained for ARID3a with goat anti-human ARID3a peptide-specific antibody, as described previously ^12^. Donkey anti-goat IgG FITC (sc-2024; Santa Cruz Biotechnology). Data were collected on a Stratedigm S1200Ex and analysis was performed using FlowJo (Tree Star) software version 10.

### Single-cell RNA-seq

FACS-sorted naïve B cells from ten randomly selected SLE patients were immediately processed on the C1 Single-Cell Auto Prep System (Fluidigm), as we have previously published ^18^. Cells were loaded onto a 5-10 µm RNA-seq microfluidic IFC plate (Fluidigm Cat# 100-5759) and capture site occupancy was confirmed microscopically. The SMARTer Ultra Low RNA Kit v4 (Clontech Cat# 635026) for full-length smart-seq was used for lysis, reverse transcription, and cDNA synthesis. Illumina sequencing libraries were constructed using the Nextera XT kit (Illumina Cat# FC 131-1096). Sequencing library quality was determined by analyzing fragment size distribution using the Agilent High Sensitivity D1000 kit on an Agilent 2200 TapeStation (Agilent Technologies) at the Oklahoma Medical Research Foundation Genomics Core Facility. Paired-end (2 x 50 bp) sequencing was performed on a NovaSeq 6000 platform and paired-end reads were aligned to the UCSC hg38 transcriptome ^38^. These alignments were used as input in RSEM v1.3 to quantify gene expression levels (transcripts per million, TPM) for all UCSC hg38 genes in all samples. A total of 901 cells were sequenced. After removal of cells of questionable viability (> 25% mitochondrial gene reads and <50 expressed genes), 829 cells were used for differential expression analyses based on levels of ARID3a transcript expression. All genes not expressed at appreciable levels ((log2 + 1) > 1) in at least 1% of single cells were discarded, leaving 8,491 total genes analyzed. Cells with greater than 25% of counts mapping to mitochondrial genes were discarded as stressed or dying cells and eliminated from downstream analyses, leaving a total of 500 cells for Seurat ^39^ analyses for unsupervised hierarchical clustering. Data were analyzed Log normalized expression values were used for uniform manifold clustering and differential expression among the clusters. The Partek Genomics Suite ComBat program ^40^ was used to remove batch effects and single cell data was segregated based on ARID3a expression, with > 0.5 log_2_ TPM denoting ARID3a^+^ cells and < 0.5 log_2_ TPM denoting ARID3a^-^ cells. ANOVA analysis was performed within the Partek Flow software to identify differentially expressed genes (DEGs) with a fold-change ≥ ± 2 and a false discovery rate (FDR) < 0.05.

### Analyses

Paired flow cytometry data was evaluated for statistical significance (Fig 6C, D; GraphPad Prism 10.5.0). Continuous variables were first assessed for normality (Shapiro-Wilk test). All differences were normally distributed (p>0.1), and the paired t test was used to compare B cells cultured with or without ARID3A-specific morpholino.

## Acknowledgements

The authors would like to thank Dr. Pat Gaffney for helpful discussions, Dr. Mary Beth Humphrey for reviewing the manuscript and the Flow Cytometry Core Facilities at OMRF, and at OU which was subsidized by the Stephenson Cancer Center.

## Funding

This work was supported by R56-AI174413 and DOD HT9425-23-1-0271, LR220036 (CFW).

## Author contributions

CFW conceived and designed the study. CFW and JG designed experiments and wrote the manuscript. CFW, JG, JMG and JH performed experiments. JG, LG, HZ and KZ contributed to scRNA-seq analyses and statistics. JJ provided patient samples and helpful discussions. All authors analyzed data and reviewed the manuscript.

## Competing interests

A patent application based on the findings of these studies has been filed by the University of Oklahoma Health Center. The authors declare that they have no other competing interests.

## Data availability

RNA sequencing data sets are available from the National Library of Medicine at GEO GSE337150.

## Supplementary Figure Legends

**Table S1.** Antibodies Used.

| Antibody | Amount | Vendor | Catalog# |
| --- | --- | --- | --- |
| **Flow cytometry** |  |  |  |
| IgD-PerCP-Cy5.5 | 1:50 | BioLegend | 348208 |
| IgM-PacBlue | 1:100 | BioLegend | 314516 |
| CD27-PE | 1:100 | BioLegend | 311106 |
| CD27-BV785 | 1:100 | BioLegend | 302832 |
| CD10-APC | 1:100 | BioLegend | 312210 |
| CD20-APCFire750 | 1:200 | BioLegend | 302358 |
| CD3-BV421 | 1:200 | BioLegend | 317344 |
| CD24-BV785 | 1:100 | BioLegend | 302832 |
| IgD-PerCP/Cy5.5 | 1:50 | BioLegend | 307810 |
| IgM-BV605 | 1:100 | BioLegend | 314524 |
| CD19-APCFire750 | 1:100 | BioLegend | 363030 |
| CD21-PE-Dazzle-594 | 1:50 | BioLegend | 354922 |
| CD138-BV421 | 1:50 | BioLegend | 356516 |
| CD68-PE | 1:20 | BioLegend | 333808 |
| CD11c-APC | 1:50 | BD Pharmingen | 559877 |
| CxCR5-APC | 1:100 | R&D Systems | FAB190A |
| CD38-BV421 | 1:100 | BioLegend | 303526 |
| CD36-PE | 1:100 | BioLegend | 336206 |
| HLA-DR-BV510 | 1:100 | BD Bioscience | 563083 |
| fixation buffer |  | BioLegend | 420801 |
| permeabilized with<br>FoxP3/Transcription Factor Staining<br>Buffer Set |  | Invitrogen<br>eBioscience | 00552300 |
| Donkey anti-goat IgG FITC |  | Santa Cruz<br>Biotechnology | sc-2024 |
| **Cell enrichment** |  |  |  |
| Easysep human B cell enrichment kit | kit | Stemcell<br>Technologies | 19054 |
| REAlase CD19 MicroBead Kit |  | Miltenyi Biotec | 130-117-034 |
| **Cell Stimulation** |  |  |  |
| R848 | 1 µg/ml |  |  |
| F(ab')<sub>2</sub> anti-IgM | 10 µg/ml | Invitrogen | A24490 |
| BAFF | 10 ng/ml | BioLegend | 559602 |
| IL21 | 10 ng/ml | BioLegend | 571202 |
| IL-2 | 50 U/ml | BioLegend | 589102 |
| IFNγ | 20 ng/ml | BioLegend | 570204 |

**Supplementary Figure 1.**
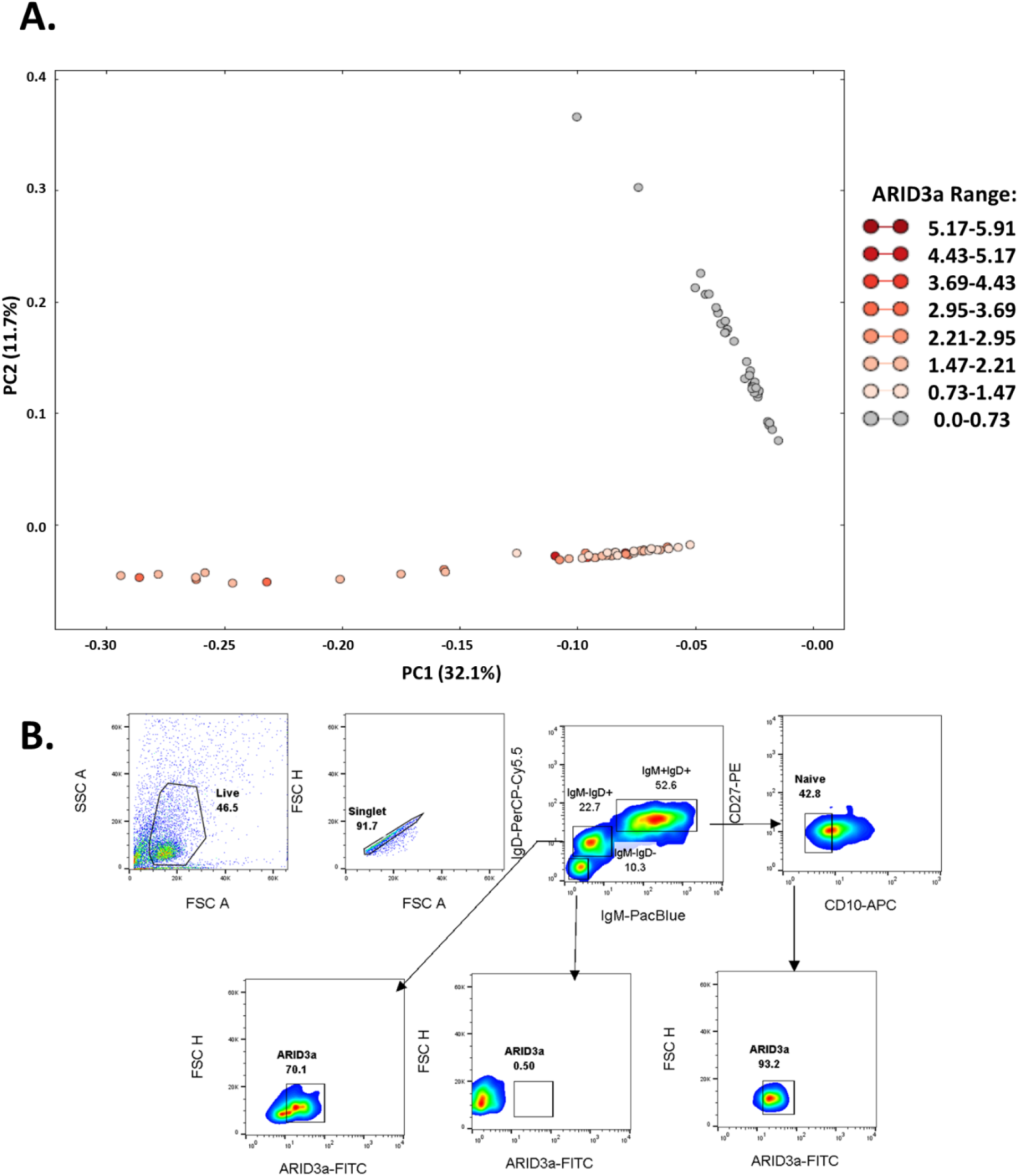
ARID3a-expressing B cells exhibit different transcriptomes from ARID3a negative B cells. A. Principal component analyses of 96 total CD19^+^ B cells from a representative SLE patient show clustering of ARID3a^+^ (red) versus ARID3a^-^ (grey) cells. B. Representative flow cytometric gating strategy for isolation of naïve B cells (IgM^+^IgD^hi^CD10^-^) shows ARID3a expression within the naïve B cell population. In this sample 84% of the cells had ARID3a transcripts.

## Notes

### Competing Interest Statement

The authors have declared no competing interest.

## References

1. Jenks, S.A., Cashman, K.S., Zumaquero, E., Marigorta, U.M., Patel, A.V., Wang, X., Tomar, D., Woodruff, M.C., Simon, Z., Bugrovsky, R., et al. (2018). Distinct Effector B Cells Induced by Unregulated Toll-like Receptor 7 Contribute to Pathogenic Responses in Systemic Lupus Erythematosus. Immunity 49, 725–739 e726. 10.1016/j.immuni.2018.08.015.

2. Tipton, C.M., Fucile, C.F., Darce, J., Chida, A., Ichikawa, T., Gregoretti, I., Schieferl, S., Hom, J., Jenks, S., Feldman, R.J., et al. (2015). Diversity, cellular origin and autoreactivity of antibody-secreting cell population expansions in acute systemic lupus erythematosus. Nat Immunol 16, 755–765. 10.1038/ni.3175.

3. Lu, R., Guthridge, J.M., Chen, H., Bourn, R.L., Kamp, S., Munroe, M.E., Macwana, S.R., Bean, K., Sridharan, S., Merrill, J.T., and James, J.A. (2019). Immunologic findings precede rapid lupus flare after transient steroid therapy. Sci Rep 9, 8590. 10.1038/s41598-019-45135-w.

4. Scharer, C.D., Blalock, E.L., Barwick, B.G., Haines, R.R., Wei, C., Sanz, I., and Boss, J.M. (2016). ATAC-seq on biobanked specimens defines a unique chromatin accessibility structure in naive SLE B cells. Sci Rep 6, 27030. 10.1038/srep27030.

5. Webb, C.F., Das, C., Coffman, R.L., and Tucker, P.W. (1989). Induction of immunoglobulin mu mRNA in a B cell transfectant stimulated with interleukin-5 and a T-dependent antigen. J Immunol 143, 3934–3939.

6. Popowski, M., Templeton, T.D., Lee, B.K., Rhee, C., Li, H., Miner, C., Dekker, J.D., Orlanski, S., Bergman, Y., Iyer, V.R., et al. (2014). Bright/Arid3A acts as a barrier to somatic cell reprogramming through direct regulation of Oct4, Sox2, and Nanog. Stem Cell Reports 2, 26–35. 10.1016/j.stemcr.2013.12.002.

7. Garton, J., Barron, M.D., Ratliff, M.L., and Webb, C.F. (2019). New Frontiers: ARID3a in SLE. Cells 8. 10.3390/cells8101136.

8. Garton, J., Shankar, M., Chapman, B., Rose, K., Gaffney, P.M., and Webb, C.F. (2021). Deficiencies in the DNA Binding Protein ARID3a Alter Chromatin Structures Important for Early Human Erythropoiesis. Immunohorizons 5, 802–817. 10.4049/immunohorizons.2100083.

9. Herrscher, R.F., Kaplan, M.H., Lelsz, D.L., Das, C., Scheuermann, R., and Tucker, P.W. (1995). The immunoglobulin heavy-chain matrix-associating regions are bound by Bright: a B cell-specific trans-activator that describes a new DNA-binding protein family. Genes Dev. 9, 3067–3082.

10. Shankar, M., Nixon, J.C., Maier, S., Workman, J., Farris, A.D., and Webb, C.F. (2007). Anti-nuclear antibody production and autoimmunity in transgenic mice that overexpress the transcription factor Bright. J Immunol 178, 2996–3006.

11. Oldham, A.L., Miner, C.A., Wang, H.C., and Webb, C.F. (2011). The transcription factor Bright plays a role in marginal zone B lymphocyte development and autoantibody production. Mol.Immunol. 49, 367–379. S0161-5890(11)00748-6 [pii];10.1016/j.molimm.2011.09.008 [doi].

12. Ward, J.M., Rose, K., Montgomery, C., Adrianto, I., James, J.A., Merrill, J.T., and Webb, C.F. (2014). Disease activity in systemic lupus erythematosus correlates with expression of the transcription factor AT-rich-interactive domain 3A. Arthritis Rheumatol. 66, 3404–3412. 10.1002/art.38857 [doi].

13. Borestrom, C., Forsman, A., Ruetschi, U., and Rymo, L. (2012). E2F1, ARID3A/Bright and Oct-2 factors bind to the Epstein-Barr virus C promoter, EBNA1 and oriP, participating in long-distance promoter-enhancer interactions. J.Gen.Virol. 93, 1065–1075. vir.0.038752-0 [pii];10.1099/vir.0.038752-0 [doi].

14. Jog, N.R., and James, J.A. (2020). Epstein Barr Virus and Autoimmune Responses in Systemic Lupus Erythematosus. Front Immunol 11, 623944. 10.3389/fimmu.2020.623944.

15. Wood, R.A., Guthridge, L., Thurmond, E., Guthridge, C.J., Kheir, J.M., Bourn, R.L., Wagner, C.A., Chen, H., DeJager, W., Macwana, S.R., et al. (2021). Serologic markers of Epstein-Barr virus reactivation are associated with increased disease activity, inflammation, and interferon pathway activation in patients with systemic lupus erythematosus. J Transl Autoimmun 4, 100117. 10.1016/j.jtauto.2021.100117.

16. Laurynenka, V., Ding, L., Kaufman, K.M., James, J.A., and Harley, J.B. (2022). A High Prevalence of Anti-EBNA1 Heteroantibodies in Systemic Lupus Erythematosus (SLE) Supports Anti-EBNA1 as an Origin for SLE Autoantibodies. Front Immunol 13, 830993. 10.3389/fimmu.2022.830993.

17. Ratliff, M.L., Garton, J., Garman, L., Barron, M.D., Georgescu, C., White, K.A., Chakravarty, E., Wren, J.D., Montgomery, C.G., James, J.A., and Webb, C.F. (2019). ARID3a gene profiles are strongly associated with human interferon alpha production. J Autoimmun 96, 158–167. 10.1016/j.jaut.2018.09.013.

18. Ratliff, M.L., Garton, J., James, J.A., and Webb, C.F. (2020). ARID3a expression in human hematopoietic stem cells is associated with distinct gene patterns in aged individuals. Immun Ageing 17, 24. 10.1186/s12979-020-00198-6.

19. Palanichamy, A., Barnard, J., Zheng, B., Owen, T., Quach, T., Wei, C., Looney, R.J., Sanz, I., and Anolik, J.H. (2009). Novel human transitional B cell populations revealed by B cell depletion therapy. J Immunol. 182, 5982–5993.

20. Nixon, J.C., Ferrell, S., Miner, C., Oldham, A.L., Hochgeschwender, U., and Webb, C.F. (2008). Transgenic mice expressing dominant-negative Bright exhibit defects in B1 B cells. J Immunol. 181, 6913–6922.

21. van der Vlist, M., Ramos, M.I.P., van den Hoogen, L.L., Hiddingh, S., Timmerman, L.M., de Hond, T.A.P., Kaan, E.D., van der Kroef, M., Lebbink, R.J., Peters, F.M.A., et al. (2021). Signaling by the inhibitory receptor CD200R is rewired by type I interferon. Sci Signal 14, eabb4324. 10.1126/scisignal.abb4324.

22. Ward, J.M., Ratliff, M.L., Dozmorov, M.G., Wiley, G., Guthridge, J.M., Gaffney, P.M., James, J.A., and Webb, C.F. (2016). Human effector B lymphocytes express ARID3a and secrete interferon alpha. J Autoimmun 75, 130–140. 10.1016/j.jaut.2016.08.003.

23. Ratliff, M.L., Shankar, M., Guthridge, J.M., James, J.A., and Webb, C.F. (2020). TLR engagement induces ARID3a in human blood hematopoietic progenitors and modulates IFNalpha production. Cell Immunol 357, 104201. 10.1016/j.cellimm.2020.104201.

24. Moulton, J.D. (2017). Using Morpholinos to Control Gene Expression. Curr Protoc Nucleic Acid Chem 68, 4 30 31-34 30 29. 10.1002/cpnc.21.

25. Gottfried, E., Kunz-Schughart, L.A., Weber, A., Rehli, M., Peuker, A., Muller, A., Kastenberger, M., Brockhoff, G., Andreesen, R., and Kreutz, M. (2008). Expression of CD68 in non-myeloid cell types. Scand J Immunol 67, 453–463. 10.1111/j.1365-3083.2008.02091.x.

26. De Groof, A., Hemon, P., Mignen, O., Pers, J.O., Wakeland, E.K., Renaudineau, Y., and Lauwerys, B.R. (2017). Dysregulated Lymphoid Cell Populations in Mouse Models of Systemic Lupus Erythematosus. Clin Rev Allergy Immunol 53, 181–197. 10.1007/s12016-017-8605-8.

27. Zhang, P., Li, W., Wang, Y., Hou, L., Xing, Y., Qin, H., Wang, J., Liang, Y., and Han, H. (2006). Identification of CD36 as a new surface marker of marginal zone B cells by transcriptomic analysis. Mol.Immunol.

28. Hardy, R.R. (2006). B-1 B cells: development, selection, natural autoantibody and leukemia. Curr.Opin.Immunol. 18, 547–555.

29. Hayakawa, K., Li, Y.-S., Shinton, S.A., Bandi, S.R., Formica, A.M., Brill-Dashoff, J., and Hardy, R.R. (2019). Crucial role of increased Arid3a at the pre-B and immature B cell stages for B1a cell generation. Frontiers in immunology 10, 457.

30. Clark, E.A., and Giltiay, N.V. (2018). CD22: A Regulator of Innate and Adaptive B Cell Responses and Autoimmunity. Front Immunol 9, 2235. 10.3389/fimmu.2018.02235.

31. Tse, K.W., Dang-Lawson, M., Lee, R.L., Vong, D., Bulic, A., Buckbinder, L., and Gold, M.R. (2009). B cell receptor-induced phosphorylation of Pyk2 and focal adhesion kinase involves integrins and the Rap GTPases and is required for B cell spreading. J Biol Chem 284, 22865–22877. 10.1074/jbc.M109.013169.

32. James, J.A., and Robertson, J.M. (2012). Lupus and Epstein-Barr. Curr.Opin.Rheumatol. 24, 383–388. 10.1097/BOR.0b013e3283535801 [doi].

33. Lanz, T.V., Brewer, R.C., Ho, P.P., Moon, J.S., Jude, K.M., Fernandez, D., Fernandes, R.A., Gomez, A.M., Nadj, G.S., Bartley, C.M., et al. (2022). Clonally expanded B cells in multiple sclerosis bind EBV EBNA1 and GlialCAM. Nature 603, 321–327. 10.1038/s41586-022-04432-7.

34. Isnardi, I., Ng, Y.S., Menard, L., Meyers, G., Saadoun, D., Srdanovic, I., Samuels, J., Berman, J., Buckner, J.H., Cunningham-Rundles, C., and Meffre, E. (2010). Complement receptor 2/CD21-human naive B cells contain mostly autoreactive unresponsive clones. Blood 115, 5026–5036.

35. Quach, T.D., Manjarrez-Orduno, N., Adlowitz, D.G., Silver, L., Yang, H., Wei, C., Milner, E.C., and Sanz, I. (2011). Anergic responses characterize a large fraction of human autoreactive naive B cells expressing low levels of surface IgM. J Immunol 186, 4640–4648. 10.4049/jimmunol.1001946.

36. Yurasov, S., Wardemann, H., Hammersen, J., Tsuiji, M., Meffre, E., Pascual, V., and Nussenzweig, M.C. (2005). Defective B cell tolerance checkpoints in systemic lupus erythematosus. J.Exp.Med. 201, 703–711.

37. Dorner, T., and Lipsky, P.E. (2024). The essential roles of memory B cells in the pathogenesis of systemic lupus erythematosus. Nat Rev Rheumatol 20, 770–782. 10.1038/s41584-024-01179-5.

38. Fujita, P.A., Rhead, B., Zweig, A.S., Hinrichs, A.S., Karolchik, D., Cline, M.S., Goldman, M., Barber, G.P., Clawson, H., Coelho, A., et al. (2011). The UCSC Genome Browser database: update 2011. Nucleic Acids Res 39, D876–882. 10.1093/nar/gkq963.

39. Butler, A., Hoffman, P., Smibert, P., Papalexi, E., and Satija, R. (2018). Integrating single-cell transcriptomic data across different conditions, technologies, and species. Nat Biotechnol 36, 411–420. 10.1038/nbt.4096.

40. Johnson, W.E., Li, C., and Rabinovic, A. (2007). Adjusting batch effects in microarray expression data using empirical Bayes methods. Biostatistics 8, 118–127. 10.1093/biostatistics/kxj037.

